# Social immunity and chemical communication in the honeybee: immune-challenged bees enter enforced or self-imposed exile

**DOI:** 10.1101/2020.08.21.262113

**Authors:** Tarli E. Conroy, Luke Holman

**Affiliations:** School of BioSciences, University of Melbourne, Royal Parade, Parkville, VIC 3010, Australia; School of Applied Sciences, Edinburgh Napier University, Edinburgh, EH11 4BN, United Kingdom

**Keywords:** Cuticular Hydrocarbons, Evolutionary immunology, Social immunity, Hygienic behaviour, Lipopolysaccharide

## Abstract

Animals living in large colonies are especially vulnerable to infectious pathogens, and may therefore have evolved additional defences. Eusocial insects supplement their physiological immune systems with ‘social immunity’, a set of adaptations that impedes the entrance, establishment, and spread of pathogens in the colony. We here find that honey bee workers (*Apis mellifera*) that had been experimentally immune-challenged with lipopolysaccharide (LPS) often exited the hive and subsequently died; some individuals were dragged out by other workers, while others appeared to leave voluntarily. In a second experiment, we found that healthy workers treated with surface chemicals from LPS-treated bees were evicted from the hive more often than controls, indicating that immune-challenged bees produce chemical cues or signals that elicit their eviction. Thirdly, we observed pairs of bees in the lab, and found that pairs spent more time apart when one member of the pair had received LPS, relative to controls. Our findings suggest that immune-challenged bees altruistically banish themselves, and that workers evict sick individuals which they identify using olfactory cues, putatively because of (kin) selection to limit the spread of pathogens within colonies.

## Introduction

Colony-living animals face a heightened risk of infectious disease, which can spread rapidly when conspecifics are in frequent contact (Naug and Camazine, 2002; Pie et al., 2004). One might therefore predict highly social species to have evolved stronger immune systems than their more solitary relatives. This prediction has received only partial support in the eusocial Hymenoptera, and indeed some evidence suggests that eusocial species have comparatively weak immune systems. Interspecific comparative analyses have found both positive and negative correlations between colony size and physiological immune responses (Stow et al., 2007; Turnbull et al., 2011; López-Uribe et al., 2016), suggesting that advanced sociality does not necessarily create selection for stronger immunity, and might even select for weaker immunity.

Inspired by the lack of a clear strengthening of the immune system among eusocial taxa, as well as observations of their behaviour, researchers have proposed that eusocial insects combat pathogens using ‘social immunity’, which provides a complementary defence system that reduces pathogen exposure and thereby weakens selection for stronger individual immune defences (Cremer et al., 2007). Social immunity is defined as the set of behavioural, physiological, and organisational adaptations that impede the entrance, establishment, and spread of pathogens in the colony (Cremer et al., 2007). For example, many eusocial insects collectively remove waste from the nest (Bot et al., 2001), or coat its interior with antimicrobial substances collected from plants or produced in their own bodies (Christe et al., 2003; Simone-Finstrom and Spivak, 2010). Many species have compartmentalised nests that help to contain the spread of pathogens; for example, leaf cutter ant nests have a separate garbage dump, and workers from the dump tend not to venture into the rest of the colony (Hart and Ratnieks, 2001). Some bees deal with infestations of parasitic mites by identifying and removing infected larvae and pupae (Spivak, 1996), or by ‘mummifying’ foreign objects or even live intruders with wax and propolis (Greco et al., 2010). Asian honey bee larvae (*Apis cerana*) were found to die more quickly upon exposure to parasitic mites than a non-coevolved species of honey bee (*A. mellifera*), leading the authors to hypothesise that *A. cerana* has evolved ‘social apoptosis’ in order to limit within-colony parasite transmission (Page et al., 2016). Pull et al. (2018) observed ant workers identifying infected ant pupae via chemical cues, and then spraying them with antimicrobial poison, in a similar manner to how diseased cells are removed by the immune system in the bodies of multicellular organisms.

Studies have also reported that honeybees show behavioural responses towards sick individuals, in ways that suggest social immunity. Baracchi *et al*. (2012) introduced newly-emerged worker bees infected with deformed wing virus (DWV) to the hive alongside healthy controls, and observed that the infected bees were frequently ejected from the hive by other workers, while controls were not. Moreover, DWV-infected bees produced a different blend of cuticular hydrocarbons, as measured by gas chromatography (Baracchi et al., 2012). Cuticular hydrocarbons (CHCs) are a layer of waxy chemicals on the body surface that prevents desiccation and has many important functions in chemical communication, e.g. in the identification of nestmates (van Zweden and d’Ettorre, 2010), the queen (Holman, 2018), and workers of differ ages and task specialties (Greene and Gordon, 2003). Secondly, Richard *et al*. (2008) found that applying chemical cues extracted from the body surface of experimentally immune-challenged bees to healthy bees caused the latter to elicit more antennation and allogrooming from nestmates. Together, these results suggest that workers can detect sick nestmates (possibly via chemical cues such as CHCs), and that they sometimes respond behaviourally by investigating, avoiding, and/or ejecting sick individuals.

Furthermore, kin selection theory (Hamilton, 1964) predicts that some social insects might have evolved ‘altruistic’ responses to sickness. In advanced eusocial species like honeybees, workers rarely breed, and instead reproduce their alleles indirectly by providing assistance to relatives such as their mother queen and her offspring. Thus, as soon as the worker’s presence flips from having a beneficial to a detrimental effect on the fitness of its relatives – such as when the worker picks up an infectious pathogen – leaving the colony permanently may become the course of action that maximises the worker’s inclusive fitness. Consistent with this idea, Rueppell *et al*. (2010) observed that worker bees exposed to harmful doses of CO_2_ or hydroxyurea flew out of the colony and did not return; the authors hypothesised this behaviour to be ‘altruistic suicide’ that evolved via kin selection to prevent harm to related nestmates. Similarly, Heinze and Walter (2010) found that ants affected by a fungal infection left their nests and remained outside until death. Another study reported that experimentally immune-challenged bees showed reduced movement, and also reduced social interactions, which was hypothesised to be an adaptation that limits disease transmission (Kazlauskas et al., 2016).

In light of this research, we hypothesise that *Apis mellifera* honeybees (and perhaps other eusocial insects) use a multi-pronged approach to social immunity that involves both collective (e.g. eviction of sick individuals by the society) as well as individual behaviours (such as altruistic self-isolation). We here investigate these ideas using behavioural experiments on social immunity involving bacterial lipopolysaccharides (LPS), which are non-living cell wall components found in Gram-negative bacteria. LPS elicits a strong response from the innate immune system in many organisms, including honey bees (Imler et al., 2000; Aubert and Richard, 2008; Kazlauskas et al., 2016). LPS is experimentally useful because it is non-living and thus cannot manipulate host behaviour as some pathogens and parasites do (Hughes et al., 2012); this is important because social immunity is about adaptive behaviours in hosts (not pathogens). In Experiment 1, we treated worker bees with LPS or two different procedural controls, re-introduced them to their natal hive, and then estimated the rates at which bees 1) remained inside, 2) left voluntarily, or 3) were ejected by other workers. In Experiment 2, we transferred surface chemicals from immune-challenged bees to healthy bees, and tested whether the healthy bees were evicted from the hive more often than controls. In Experiment 3, we observed pairs of bees in which one member was immune-challenged, to test whether they became more or less gregarious relative to controls.

## Methods

### Experimental animals

We utilised five honeybee colonies, each housed in a Langstroth hive on the Parkville campus of The University of Melbourne. Most of the work was carried out in in April – June 2019, except one block of Experiment 1 conducted in December 2019. The workers used in our experiments were collected by opening up a hive and selecting a centrally-located frame containing developing brood, then brushing a haphazardly-selected sample of workers clinging to the frame into a plastic container for transport to the lab. By selecting within-nest bees that did not fly in response to the disturbance of opening the hive, we aimed to preferentially collect younger bees that have not yet begun performing outside-nest tasks such as guarding and foraging. Therefore, our default expectation is that most of these bees would remain inside when returned to the hive.

### Experimental immune challenge and controls

Following similar experiments (e.g. Kazlauskas et al., 2016), we diluted LPS (from serotype 055:B5 *E. coli*; Sigma-Aldrich) to 0.5mg/mL in a sterile physiological saline (Ringer’s solution, prepared from autoclaved, double-distilled water), and then stored it in aliquots at –18°C prior to the experiments. For all three experiments, we used Ringer’s solution with no added LPS (henceforth ‘Ringers’) as a control for the process of wounding bees with saline; aliquots of Ringers were prepared and frozen at the same time as the LPS-containing aliquots, from the same batch of Ringer’s solution.

To administer the Ringers and LPS solutions, bees were anaesthetised in small groups (30-40 individuals) by placing them in a –18°C freezer inside a plastic container until they were immobile but still moving their appendages (typically *c*. 6 minutes). We then kept the containers of bees over ice, and monitored them to maintain this state of light anaesthesia. Using a stereomicroscope and an entomological pin (0.25mm; sterilised in ethanol and a candle flame between uses), we then randomly selected a bee, dipped the pin into one of the treatment solutions, and then inserted the pin through the pleural membrane between the fourth and fifth tergal segments to a distance of roughly 1mm (using a different pin for each treatment solution). Bees in Experiment 1’s ‘Intact control’ group were handled similarly (i.e. anaesthetised and manipulated under the microscope), but were not punctured with a pin. After treatment, bees were marked on the thorax with a dot of coloured paint to identify their treatment group; we used a different pairing of colours and treatments for each experimental replicate to prevent confounding. For all experiments, we applied the treatments on a rotation, preventing confounding effects from e.g. order of processing and time under anaesthesia.

#### Experiment 1: Do immune-challenged bees leave the hive?

Experiment 1 utilised three treatments: an intact control, and treatments in which bees were punctured with a needle coated in Ringers or LPS. After applying these treatments as described, bees were housed in their treatment groups, in the dark at 25°C, for 18±1 hours. We then removed any bees that had died or showed impaired mobility, and reintroduced the remainder to the hive, by opening the hive and returning them to the central frame from where they had been collected. The hive entrance was then recorded for up to two hours by an observer who stood by the hive (mean observation time: 97.5 minutes). We also video recorded the hive entrance to double-check each observation (done blind by a second observer).

We recorded a categorical response variable with three possible outcomes: bees either stayed inside the hive for the duration of the observation period, left the hive voluntarily, or were forced out. We recorded a bee as leaving voluntarily when it walked or flew out of the hive without interacting with any other workers (in practice, only 2 bees left by flying; the other 22 walked out). We recorded the ‘forced out’ outcome when the focal bee was pulled out of the hive by one or more other workers using their mandibles and thrown/dropped from the entrance of the hive. Only bees that left the landing board at the hive entrance were classified as having exited the hive. Four bees emerged from the hive and then re-entered (1× intact control, 1× Ringers, 2× LPS); the meaning of this behaviour is not clear, so these four observations were excluded from analysis. Experiment 1 was replicated over four hives: three in Autumn (30/04, 3/05 and 9/05 in 2019), and one in Spring/Summer (19/11/2019; added to increase sample size).

#### Experiment 2: The role of cuticular odours

Experiment 2 involved two treatments. We first collected a sample of *c*. 200 workers from inside a hive, anaesthetised them, and punctured them with either Ringers or LPS in Ringers (as in Experiment 1, except that we did not apply a paint mark). We then housed these bees at 25°C in the dark for 24 hours in their treatment groups, to give time for changes in the cuticular odour profile to occur in response to the experimental and control manipulations. Earlier experiments have shown that 24 hours is sufficient for LPS-treated insects to develop a substantially different chemical profile relative to Ringers-treated controls (Holman et al., 2010); indeed, Richard *et al*. (2008) observed changes after 4 hours.

Next, we freeze-killed the surviving bees and placed an equal number from each treatment into two 20mL glass vials. We then added 500*µ*L hexane (HPLC grade; Sigma ref. 34859) per bee, and then gently shook the vials by hand for 60 seconds to facilitate dissolution of hexane-soluble epicuticular chemicals, then pipetted the extract into aliquots in 2mL glass vials.

On the same day that we prepared the chemical extracts, we collected a further 200 bees from the same hive, which were cold-immobilised and then marked with one of two paint colours. As before, we used different treatment-colour pairings for each hive to avoid confounding. Because hexane is toxic, we applied the extracted CHCs to the bees indirectly by pipetting 20*µ*L of the appropriate hexane solution onto the surface of deionised water in a 10mL glass beaker (Smith et al., 2012). After waiting a few seconds for the hexane to evaporate, we then dipped an anaesthetised, paint-marked bee through the surface of the water, and swirled it in the water’s surface to allow it to pick up hydrophobic solutes (e.g. cuticular hydrocarbons) that were floating on the water’s surface. The odour-coated bees were then reintroduced to their hive, and their subsequent emergence was recorded over the next 1-2 hours (mean 97.5 minutes), as in Experiment 1. Experiment 2 was replicated across four hives between 26^th^ May and 28^th^ June 2019.

#### Experiment 3: ‘Social distancing’ following immune challenge

Like Experiment 2, this experiment had two treatments: Ringers and LPS. We collected approximately 250 bees from a brood frame as before, paired them at random, and then randomly assigned each pair to one of the two treatments. One of the bees in each pair was punctured with either Ringers or LPS, while the other was left intact. Each pair was placed into a 22mL glass test tube stoppered with cotton wool. All of the test tubes were put into a ZebraTower video recording cabinet (Viewpoint, France), then recorded under infrared illumination (invisible to bees). We then analysed the videos using ‘scan sampling’; specifically, we examined video stills separated by 120s intervals, and recorded whether or not each pair of bees was in close contact (defined as within 1.5cm of each other) in each video still (i.e. the response variable was binary). The observation period lasted 3.5 hours, and began 30 minutes after closing the video recording cabinet to allow time for the bees to settle following the disturbance. Videos were transcribed blind with respect to treatment and hive. Experiment 3 was replicated across four hives in the same time period as for Experiment 2.

### Statistical analysis

Our main statistical analyses for Experiments 1 and 2 use an uncommon model type – multinomial logistic models (hereafter MLM) – which we believe is the right choice because these experiments have a categorical response variable with three possible outcomes. However, because MLM is unfamiliar, we first present a similar analysis of the Experiment 1 and 2 data using a more common type of model (a binomial generalised linear mixed model; GLMM), to demonstrate that the main conclusions are not unique to the MLM. For the GLMM, we first re-coded the response variable to have only two outcomes, namely ‘Stayed inside’ or ‘Left the hive’ (i.e. the sum of the ‘Forced out’ and ‘Left voluntarily’ outcomes), then ran a GLMM with treatment as a fixed effect and hive as a random effect. GLMMs were implemented in the lme4 package for R. Note that the GLMM requires discarding information about which bees left by force *vs* voluntarily, making the MLM preferrable.

The MLM is an extension of binomial models to data with *>*2 outcomes. We implemented the MLM in a Bayesian framework via the R package brms (Bürkner, 2017). As in the GLMM, we included treatment as a fixed effect and hive as a random effect. We specified a prior distribution of N(*µ* = 0, *σ*^2^ = 3) for the fixed effect estimates and N(*µ* = 0, *σ*^2^ = 2) for the random effects, in order to ‘regularise’ the parameter estimates (see McElreath, 2020) and help model convergence.

Experiment 3 had a binary response variable and was therefore modelled using a GLMM (implemented in brms). The model included treatment, hive, and the additional random factor ‘pair ID’, to account for repeated measurements of each pair of bees; hive was fitted as a fixed factor to ensure the model converged.

We ran four chains for the Bayesian models of Experiments 1 and 2 (5000 iterations per chain; first 2500 discarded as burn-in), and confirmed model convergence and fit via 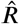 statistics and posterior predictive plots. The model for Experiment 3 required additional iterations (20,000 per chain, with 10,000 burn-in) to ensure adequate effective sampling. To make inferences from each model we ran ‘planned contrasts’, i.e. calculating the posterior estimate of the difference in means between each pair of treatments, then finding the log odds ratio (and its 95% CIs) as a measure of effect size. All data and R scripts are documented at lukeholman.github.io/social_immunity.

## Results

### Experiment 1: Do immune-challenged bees leave the hive?

In the preliminary GLMM, there was a significant effect of treatment (Wald χ^2^ test; *p* = 0.00012) on the proportion of bees leaving the hive. More bees left in the LPS treatment relative to the intact control (contrast; *p <* 0.0001), though the increase in LPS relative to the Ringers control was marginally non-significant (*p* = 0.083). Non-significantly more bees left the hive in the Ringers control compared to the intact control (*p* = 0.086).

Figure 1A uses the results of the Bayesian MLM to display the posterior estimates of the percentages of bees from each treatment group that stayed inside the hive, left voluntarily, or were forced out, while Figure 1B shows the standardised effect size for contrasts between treatment groups for each of the three response categories. Table S1 shows the MLM’s parameter estimates, and Table S2 shows the results of planned contrasts comparing each pair of treatments (as in Figure 1B).

**Figure 1:**
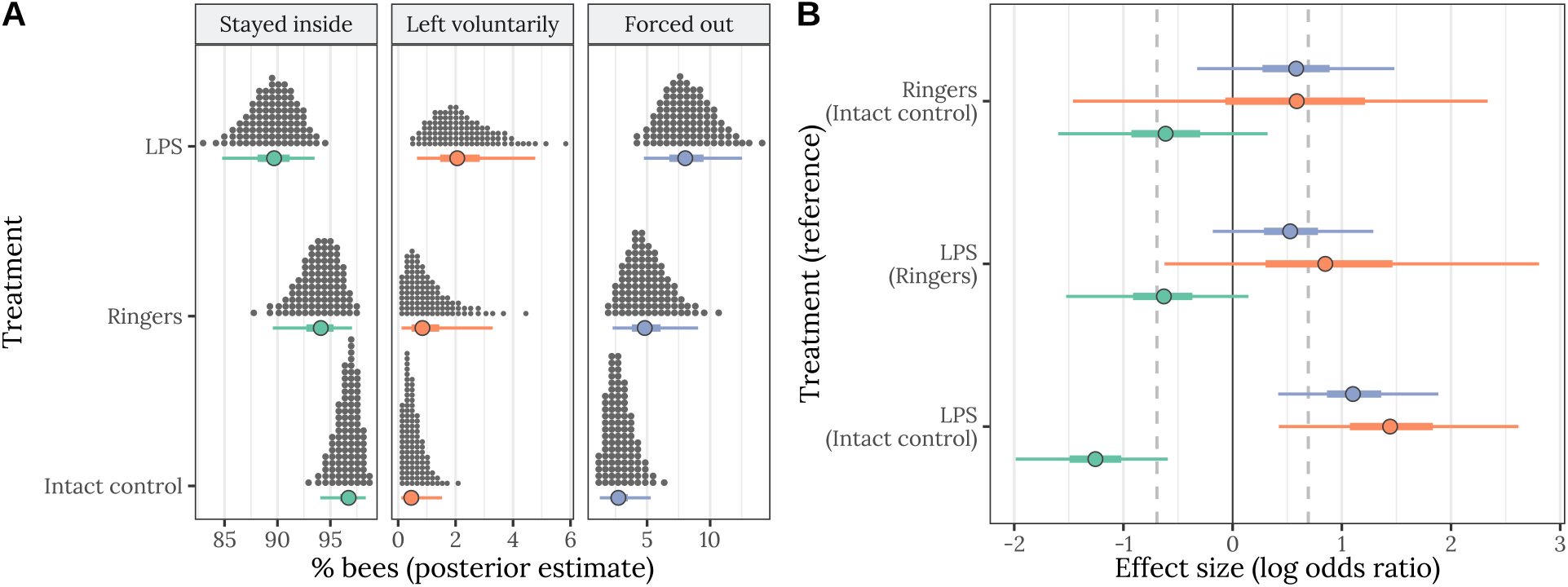
Results of Experiment 1 (n = 842 bees). Panel A shows the posterior estimate of the mean % bees staying inside the hive (left), leaving voluntarily (middle), or being forced out (right), for each of the three treatments. The quantile dot plot shows 100 approximately equally likely estimates of the true % bees, and the horizontal bars show the median and the 50% and 95% credible intervals of the posterior distribution. Panel B gives the posterior estimates of the effect size of each treatment, relative to one of the other treatments (the name of which appears in parentheses), expressed as a log odds ratio (LOR). Positive LOR indicates that the % bees showing this particular outcome is higher in the treatment than the control; for example, more bees left voluntarily (orange) or were forced out (blue) in the LPS treatment than in the intact control. The dashed lines mark *LOR* = 0, indicating no effect, and *LOR* = ±*/og*(2), i.e. the point at which the odds are twice as high in one treatment as the other.

Bees treated with LPS were significantly less likely to stay inside the hive compared to the intact control: the posterior probability that the true effect size is negative was over 99.99% (Table S2); this percentage can be interpreted similarly to a one-tailed *p*-value of 0.0001. Hereafter, we write PP to represent 1 minus this posterior probability, for easier comparison with the familiar *p*-value. Furthermore, bees that received LPS were non-significantly less likely to stay inside the hive than those that received the Ringers control (PP = 0.0516), providing weak evidence that the immune challenge from LPS caused bees to leave even more often than bees that were wounded but not otherwise immune-challenged. Bees were also somewhat more likely to leave the hive if treated with Ringers, relative to the intact control (PP = 0.072). There were corresponding, opposite treatment effects on the numbers of bees that were forced out, or left voluntarily (Figure 1, Table S2), though the differences were more apparent for the ‘Forced out’ outcome (because this outcome was more common, increasing statistical power; Figure 1A). In particular, bees that received LPS were non-significantly more likely to be forced out than those that received the Ringers control (PP = 0.074). The treatment effect size was quite large (Figure 1B, Table S2); for example, the log odds ratio comparing the frequency of the ‘Forced out’ outcome between the Intact control and LPS treatments was 1.12, indicating that LPS-treated bees were forced out *e*^1.12^ = 3.1-fold more often.

### Experiment 2: Role of cuticular odours in behavioural responses to immune challenge

In the preliminary GLMM analysis, there was a significant effect of treatment (Wald χ^2^ test; *p* = 0.0155) on the proportion of bees leaving the hive. More bees left in the group perfumed with odours from LPS-treated bees, relative to the control group perfumed with odours from Ringers-treated bees.

Figure 2A uses the results of the Bayesian MLM to display the posterior estimates of the percentages of bees from each treatment group that stayed inside the hive, left voluntarily, or were forced out, while Figure 2B shows the standardised effect size of the LPS treatment, for each of the three response categories. Table S3 shows the model’s parameter estimates, and Table S4 shows the results of planned contrasts comparing each pair of treatments.

**Figure 2:**
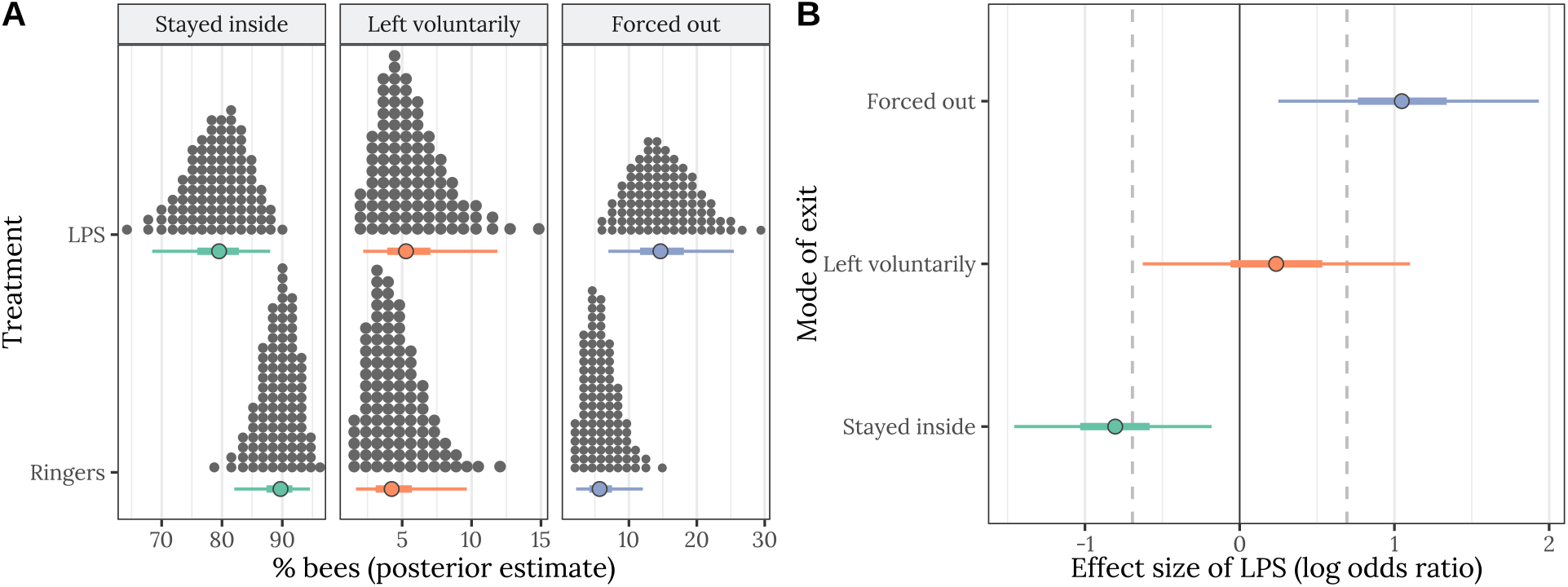
Results of Experiment 2 (n = 585 bees). Panel A shows the same information as Figure 1A. Panel B gives the posterior estimates of the effect size (log odds ratio) of the LPS treatment as a log odds ratio, for each of the three possible outcomes; the details are the same as in Figure 1B.

Figures 2A-B and Tables S3-4 illustrate that bees coated with chemical extracts from LPS-treated bees were forced out of the hive significantly more often than were those treated with chemical extracts from Ringers-treated bees (PP = 0.0046). The effect size of LPS was quite large (Log odds ratio: 1.08; Figure 2B, Table S4), indicating that LPS-perfumed bees were forced out *e*^1.08^ = 2.9-fold more often than the Ringers-perfumed controls. Interestingly, there was no significant difference in the rate at which the two treatment groups left the hive voluntarily (PP = 0.29), as expected given that the focal bees in Experiment 2 were intact and not immune-challenged.

### Experiment 3: ‘Social distancing’ following immune challenge

Figure 3 illustrates that pairs of bees in which one individual had been treated with LPS spent less time in close contact than pairs in which one individual had received Ringers. Figure 3A shows histograms of the raw data, illustrating that bees in the control group more often spent *>*90% of the 3.5 hour observation in close contact, while LPS treatment bees were over-represented among pairs that spent a lot of time apart. Tables S5-6 give the associated statistical results; the mean % time in close contact (Figure 3B) differed statistically significantly between treatments (PP = 0.032), and the effect size was moderate (log odds ratio: 0.37).

**Figure 3:**
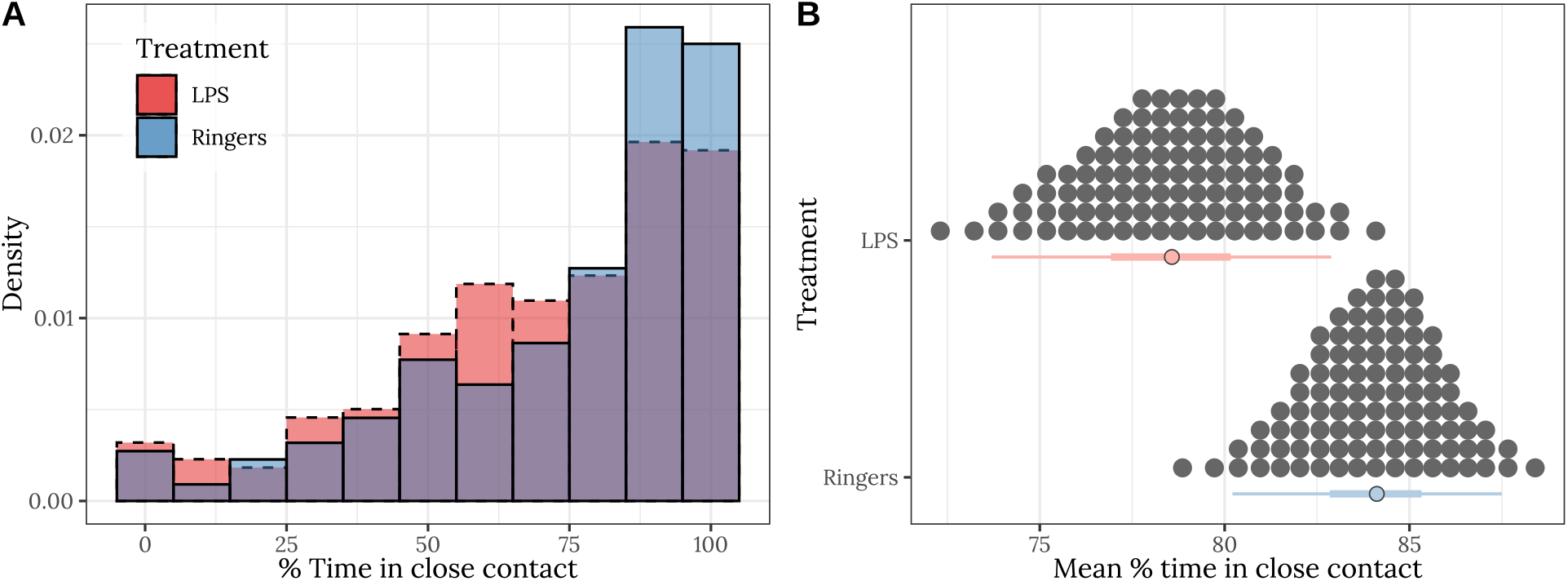
Results of Experiment 3 (n = 439 bees). Panel A shows the frequency distribution of the % time in close contact, for pairs of bees from the LPS treatment and the Ringers control. Panel B shows the posterior estimates of the mean % time spent in close contact; the details of the quantile dot plot and error bars are the same as described for Figure 1.

## Discussion

Experiment 1 revealed that bees treated with LPS were more likely to leave the hive compared to the intact control and (marginally non-significantly; *p* = 0.052) the Ringers control. Most of the bees that left the hive in Experiment 1 were forced out by other colony members, though many appeared to leave voluntarily, by walking out and then dropping to the ground without any apparent involvement from other workers (very few left by flying). In this sense, our results differ from those of an earlier study (Rueppell et al., 2010), which reported that bees with experimentally-impaired health instead left the hive by flight. This discrepancy may reflect our studies’ different methodologies; for example, the other study challenged bees with CO_2_ or hydroxyurea, rather than an LPS-coated pin. We frequently found the hive-exiting bees dead on the same or subsequent days, and many were carried off by Vespid wasps within seconds of dropping clear of the hive, illustrating that the hive-exiting behaviour is unlikely to represent an attempt to temporarily self-quarantine. From Experiment 1, we conclude that experimentally wounded, immune-stimulated bees tend to leave the hive, both with and without any apparent involvement from other workers. Therefore, our study reaffirms the existence of both the “altruistic self-removal” reported by Rueppell *et al*. (2010) and the forceful ejection of sick individuals observed by Richard *et al*. (2008) in honeybees. The relative importance of wounding (which occurred in the Ringers control and LPS treatment, but not the intact control) and immune challenge (LPS only) in generating these behavioural responses are unclear, given the borderline-significant difference between the Ringers and LPS treatments, though the ordering of the treatment means (Figure 1A) implies that both wounding and immune challenge may affect behaviour.

The removal of immune-stimulated bees by their nestmates implies that sick individuals produce signals or cues which allow them to be identified for removal from the colony. We hypothesised that at least some of these signals/cues would be olfactory, since Hymenoptera rapidly develop a distinct chemical profile after an immune challenge (Richard et al., 2008; Holman et al., 2010). We tested this hypothesis in Experiment 2, and found that bees coated with hexane-soluble chemicals extracted from the body surface of immune-challenged bees were ejected from the colony about three-fold more often than controls, which were treated with chemicals from Ringers-treated bees. Interestingly, there was no treatment effect on the rate at which bees left the hive voluntarily. We hypothesise that this difference stems from the mismatch created in Experiment 2 between the personal information held by the experimental individuals (which correctly perceived themselves to be healthy), and the chemical cues on their body surface (which came from immune-challenged individuals), causing the bees to be forced out by their nestmates, which perceived the experimental individuals to be sick.

In light of past findings that immune-challenged ants (Bos et al., 2012) and bees (Richard et al., 2008; Kazlauskas et al., 2016) engage in fewer social interactions, Experiment 3 tested whether pairs of bees in which one member had received an immune challenge spent less time in close contact than control pairs. We recorded a statistically significant decline in the proportion of time spent in contact, suggesting a behavioural response to immune challenge in the treated individual, the healthy individual paired with them, or both. A previous study recorded that bees directed more aggression and grooming behaviours towards immune-challenged bees (Richard et al., 2008); behavioural effects like this could underpin our results. Another study recorded that LPS-treated bees showed reduced locomotion and antennated other individuals less often, which might also explain our results. Such ‘sickness behaviour’ might have been shaped by kin selection to limit disease transmission (Kazlauskas et al., 2016), though there may also be direct fitness benefits (or non-adaptive explanations) for sickness behaviour, especially given that it occurs in non-eusocial animals (e.g. Sullivan et al., 2016).

The chemical cues that distinguish healthy and immune-challenged individuals remain to be determined. Cuticular hydrocarbons (CHCs) are one likely possibility, given that honeybees utilise CHCs for chemical recognition in several other contexts (e.g. van Zweden and d’Ettorre, 2010). Furthermore, the CHC profile changes rapidly following an immune challenge, in both eusocial and non-eusocial insects. For example, honeybee workers injected with Gram-negative bacteria began to produce relatively more unsaturated and shorter-chained hydrocarbons within 6h, and there were concommitant changes in the expression of genes involved in CHC biosynthesis (Richard et al., 2012). Another reason to suspect CHCs is that the insect innate immune response involves changes in the expression of genes that have pleiotropic effects on lipid metabolism/homeostasis, such as those in the IMD pathway (e.g. Pull et al., 2018; Kamareddine et al., 2018), such that there are a plausible mechanistic links between the CHC profile and immune status. However, no study has yet manipulated the CHC profile without confounds – our study and earlier, similar experiments (Richard et al., 2008) involved solvent washes rather than treatment with CHCs specifically – so the involvement of CHCs remains to be demonstrated conclusively.

Another outstanding question is whether the changes in the external chemical cues of immune-challenged bees represent an adaptation, or simply a non-adaptive consequence of other processes (i.e. a ‘spandrel’; Gould and Lewontin, 1979). Under the adaptive hypothesis, sick bees that purposefully signal their illness would be the superorganismal equivalent of infected vertebrate cells, which use MHC class I proteins to present antigens to cytotoxic T cells, which then destroy the infected cell. Presumably, the antigen-presenting system evolved adaptively (Forni et al., 2014); it could be framed as a kin-selected adaptation because the self-sacrificing cells confer a benefit to genetically identical cells in the same body. Under the non-adaptive model, immune-challenged bees might produce modified chemical cues for reasons other than eliciting their own removal, e.g. because of pleiotropic links between immunity and metabolism (Kamareddine et al., 2018); the key feature distinguishing these two hypotheses is the presence of a net inclusive fitness benefit to workers that solicit their own removal. In support of the non-adaptive hypothesis, immune challenge has also been found to affect the CHC profile of non-social insects that appear to have no need for social immunity (e.g. the beetle *Tenebrio molitor*; Nielsen and Holman, 2012). To begin establishing whether chemical signalling of immune status has evolved adaptively, one could test whether social insects undergo uniquely strong chemical or behavioural changes following an immune challenge, relative to non-social insects, in a formal phylogenetic study.

## Acknowledgements

We are grateful to Jean-Pierre Scheerlinck, Heidi Wong, and Daisy Kocher for assisting the experiments.

## Data and code availability

Raw data and documented R code and output is available at lukeholman.github.io/social_immunity.

## Online Supplementary Material

The figures and tables in this document, along with the with the R code used to generate them, can also be viewed online at https://lukeholman.github.io/social_immunity/

**Table S1:** Table summarising the posterior estimates of each fixed effect in the Bayesian multinomial logistic model of Experiment 1. Because there were three possible outcomes for each bee (Stayed inside, Left voluntarily, or Forced out), there are two parameter estimates for each predictor in the model. ‘Treatment’ is a fixed factor with three levels, and the effects shown here are expressed relative to the ‘Intact control’ group. The *p* column gives the posterior probability that the true effect size is opposite in sign to what is reported in the Estimate column, similarly to a *p*-value.

| Response | Parameter | Estimate | Est. Error | Lower<br>95% CI | Upper<br>95% CI | PP |  |
| --- | --- | --- | --- | --- | --- | --- | --- |
| % bees leaving voluntarily | Intercept | -4.94 | 0.93 | -6.97 | -3.30 | 0.0000 | *** |
|  | Treatment: Ringers | 0.59 | 0.97 | -1.43 | 2.38 | 0.2570 |  |
|  | Treatment: LPS | 1.57 | 0.56 | 0.55 | 2.73 | 0.0007 | *** |
| % bees forced out | Intercept | -3.59 | 0.81 | -5.24 | -2.00 | 0.0000 | *** |
|  | Treatment: Ringers | 0.60 | 0.46 | -0.31 | 1.50 | 0.0977 | ~ |
|  | Treatment: LPS | 1.16 | 0.37 | 0.46 | 1.93 | 0.0001 | *** |

**Table S2:**
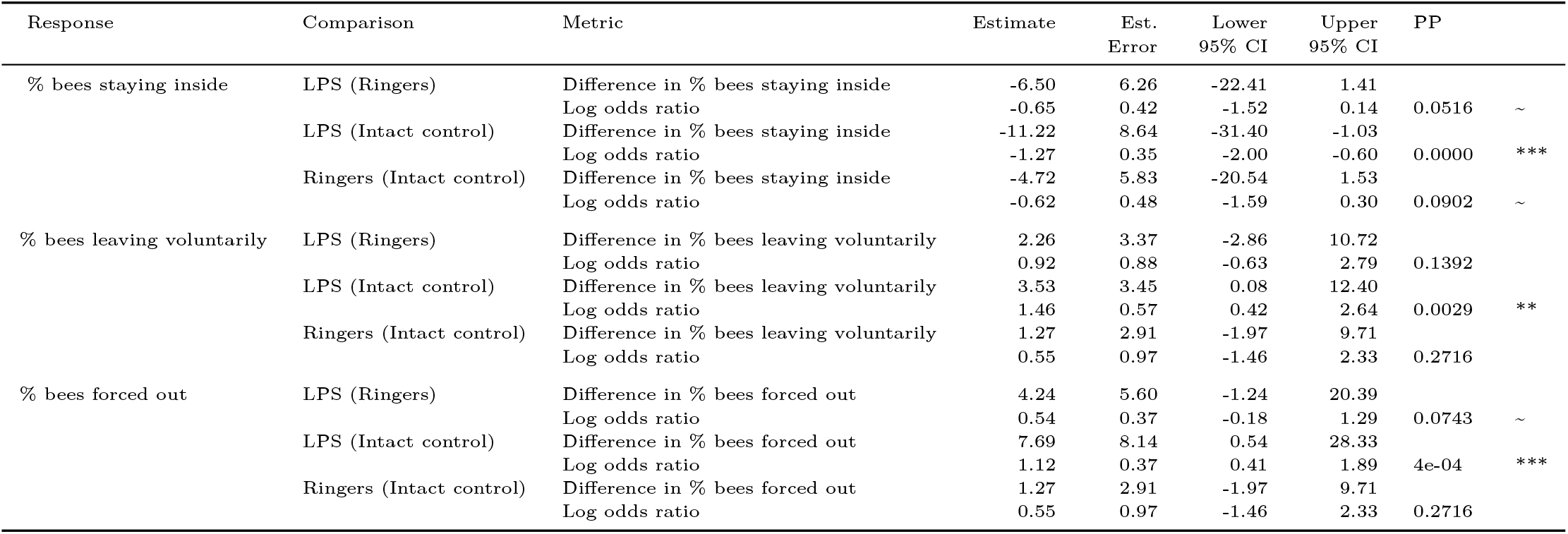
This table gives statistics associated with each of the contrasts plotted in Figure 1B. Each pair of rows gives the absolute effect size (i.e. the difference in % bees) and standardised effect size (as log odds ratio; LOR) for the focal treatment, relative to the treatment shown in parentheses, for one of the three possible outcomes (Stayed inside, Left voluntarily, or Forced out). A LOR of |*log*(*x*)| indicates that the outcome is *x* times more frequent in one treatment compared to the other, e.g. *log*(2) = 0.69 and *log*(0.5) = –0.69 correspond to a two-fold difference in frequency. The *PP* column gives the posterior probability that the true effect size has the same sign as is shown in the Estimate column; this metric has a similar interpretation to a one-tailed *p* value.

**Table S3:** Table summarising the posterior estimates of each fixed effect in the model of Experiment 2. This was a multinomial model with three possible outcomes (Stayed inside, Left voluntarily, or Forced out), and so there are two parameter estimates for each predictor in the model. ‘Treatment’ is a fixed factor with two levels, and the effect of LPS shown here is expressed relative to the ‘Ringers’ treatment. The PP column gives the posterior probability that the true effect size is opposite in sign to what is reported in the Estimate column, similarly to a *p*-value.

| Response | Parameter | Estimate | Est. Error | Lower<br>95% CI | Upper<br>95% CI | PP |  |
| --- | --- | --- | --- | --- | --- | --- | --- |
| % bees leaving voluntarily | Intercept | -3.42 | 0.55 | -4.60 | -2.39 | 0.0000 | *** |
|  | Treatment: LPS | 0.35 | 0.44 | -0.51 | 1.21 | 0.2132 |  |
| % bees forced out | Intercept | -3.65 | 0.62 | -4.96 | -2.47 | 0.0000 | *** |
|  | Treatment: LPS | 1.08 | 0.43 | 0.27 | 1.96 | 0.0046 | ** |

**Table S4:**
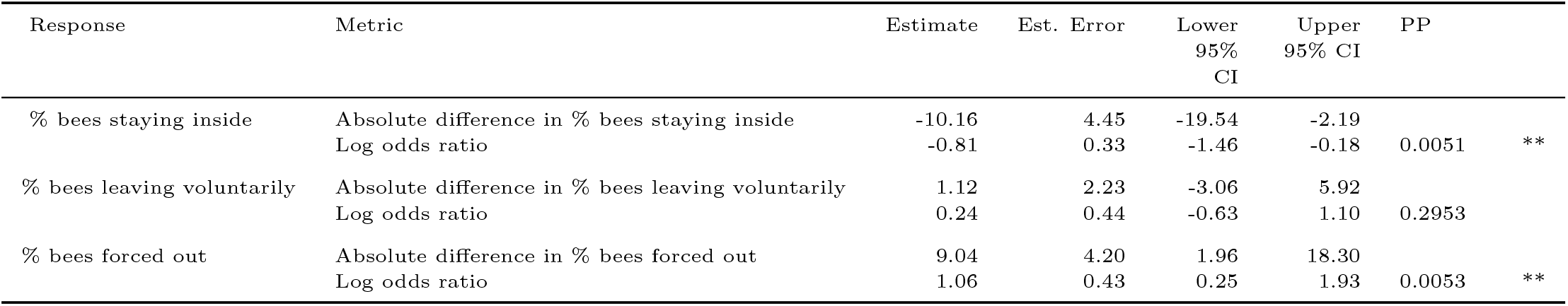
This table gives statistics associated with each of the contrasts plotted in Figure 2B. Each pair of rows gives the absolute (i.e. the difference in % bees) and standardised effect size (as log odds ratio; LOR) for the LPS treatment, relative to the Ringers control, for one of the three possible outcomes (Stayed inside, Left voluntarily, or Forced out). A LOR of |*log*(*x*)| indicates that the outcome is *x* times more frequent in one treatment compared to the other, e.g. *log*(2) = 0.69 indicates a two-fold difference in frequency. The *PP* column gives the posterior probability that the true effect size has the same sign as is shown in the Estimate column; this metric has a similar interpretation to a one-tailed *p* value in frequentist statistics.

**Table S5:** Table summarising the posterior estimates of each fixed effect in the model of Experiment 3 (a binomial GLMM where the response variable was 0 when bees were not in close contact, and 1 when they were). ‘Treatment’ is a fixed factor with two levels, and the effect of LPS shown here is expressed relative to the ‘Ringers’ treatment. ‘Hive’ was a fixed factor with four levels (modelled using deviation coding). The model also included one random effect, ‘pair ID’, which grouped observations made on each pair of bees, preventing pseudoreplication. The *PP* column gives the posterior probability that the true effect size is opposite in sign to what is reported in the Estimate column, similarly to a *p*-value.

| Parameter | Estimate | Est. Error | Lower 95% CI | Upper 95% CI | PP |  |
| --- | --- | --- | --- | --- | --- | --- |
| Intercept | 1.67 | 0.14 | 1.39 | 1.94 | 0.0000 | *** |
| Treatment: LPS | -0.37 | 0.20 | -0.75 | 0.02 | 0.0322 | * |
| hive1 | -0.20 | 0.18 | -0.55 | 0.15 | 0.1343 |  |
| hive2 | 0.14 | 0.16 | -0.17 | 0.46 | 0.1860 |  |
| hive3 | -0.23 | 0.18 | -0.58 | 0.12 | 0.0999 | ~ |

**Table S6:** Pairs in which one bee had received LPS were observed in close contact less frequently than pairs in which one bee had received Ringers. The *PP* column gives the posterior probability that the true effect size is opposite in sign to what is reported in the Estimate column, similarly to a *p*-value.

| Metric | Estimate | Est.Error | Lower 95% CI | Upper 95% CI | PP |  |
| --- | --- | --- | --- | --- | --- | --- |
| Absolute difference in % time in close contact | 5.55 | 3.0 | -0.35 | 11.45 |  |  |
| Log odds ratio | -0.37 | 0.2 | -0.75 | 0.02 | 0.0322 | * |

## References

Aubert, A., and F.-J. Richard. 2008. Social management of LPS-induced inflammation in Formica polyctena ants. Brain, Behavior, and Immunity 22:833–837.

Baracchi, D., A. Fadda, and S. Turillazzi. 2012. Evidence for antiseptic behaviour towards sick adult bees in honey bee colonies. Journal of Insect Physiology 58:1589–1596.

Bos, N., T. Lefèvre, A. Jensen, and P. d’Ettorre. 2012. Sick ants become unsociable.Journal of Evolutionary Biology 25:342–351.

Bot, A. N., C. R. Currie, A. G. Hart, and J. J. Boomsma. 2001. Waste management in leaf-cutting ants. Ethology Ecology & Evolution 13:225–237.

Bürkner, P.-C. 2017. Advanced Bayesian multilevel modeling with the R package brms. arXiv preprint:1705.11123.

Christe, P., A. Oppliger, F. Bancalà, G. Castella, and M. Chapuisat. 2003. Evidence for collective medication in ants. Ecology Letters 6:19–22.

Cremer, S., S. A. Armitage, and P. Schmid-Hempel. 2007. Social immunity. Current biology 17:R693–R702.

Forni, D., R. Cagliani, C. Tresoldi, U. Pozzoli, L. De Gioia, G. Filippi, S. Riva, G. Menozzi, M. Colleoni, M. Biasin, et al. 2014. An evolutionary analysis of antigen processing and presentation across different timescales reveals pervasive selection. PLoS Genetics 10:e1004189.

Gould, S. J., and R. C. Lewontin. 1979. The spandrels of San Marco and the Panglossian paradigm: a critique of the adaptationist programme. Proceedings of the royal society of London. Series B. Biological Sciences 205:581–598.

Greco, M. K., D. Hoffmann, A. Dollin, M. Duncan, R. Spooner-Hart, and P. Neumann. 2010. The alternative pharaoh approach: stingless bees mummify beetle parasites alive. Naturwissenschaften 97:319–323.

Greene, M. J., and D. M. Gordon. 2003. Cuticular hydrocarbons inform task decisions. Nature 423:32–32.

Hamilton, W. D. 1964. The genetical evolution of social behaviour. II. Journal of Theoretical Biology 7:17–52.

Hart, A. G., and F. L. Ratnieks. 2001. Task partitioning, division of labour and nest compartmentalisation collectively isolate hazardous waste in the leafcutting ant Atta cephalotes. Behavioral Ecology and Sociobiology 49:387–392.

Heinze, J., and B. Walter. 2010. Moribund ants leave their nests to die in social isolation. Current Biology 20:249–252.

Holman, L. 2018. Queen pheromones and reproductive division of labor: a meta-analysis. Behavioral Ecology 29:1199–1209.

Holman, L., C. G. Jørgensen, J. Nielsen, and P. d’Ettorre. 2010. Identification of an ant queen pheromone regulating worker sterility. Proceedings of the Royal Society B: Biological Sciences 277:3793–3800.

Hughes, D. P., J. Brodeur, and F. Thomas. 2012. Host manipulation by parasites. Oxford University Press.

Imler, J.-L., S. Tauszig, E. Jouanguy, C. Forestier, and J. A. Hoffmann. 2000. LPS-induced immune response in Drosophila. Journal of Endotoxin Research 6:459–462.

Kamareddine, L., W. P. Robins, C. D. Berkey, J. J. Mekalanos, and P. I. Watnick. 2018. The Drosophila immune deficiency pathway modulates enteroendocrine function and host metabolism. Cell metabolism 28:449–462.

Kazlauskas, N., M. Klappenbach, A. M. Depino, and F. F. Locatelli. 2016. Sickness behavior in honey bees. Frontiers in Physiology 7:261.

López-Uribe, M. M., W. B. Sconiers, S. D. Frank, R. R. Dunn, and D. R. Tarpy. 2016. Reduced cellular immune response in social insect lineages. Biology Letters 12:20150984.

McElreath, R. 2020. Statistical rethinking: A Bayesian course with examples in R and Stan. CRC press, London.

Naug, D., and S. Camazine. 2002. The role of colony organization on pathogen transmission in social insects. Journal of theoretical Biology 215:427–439.

Nielsen, M. L., and L. Holman. 2012. Terminal investment in multiple sexual signals: immune-challenged males produce more attractive pheromones. Functional Ecology 26:20–28.

Page, P., Z. Lin, N. Buawangpong, H. Zheng, F. Hu, P. Neumann, P. Chantawannakul, and V. Dietemann. 2016. Social apoptosis in honey bee superorganisms. Scientific Reports 6:1–6.

Pie, M. R., R. B. Rosengaus, and J. F. Traniello. 2004. Nest architecture, activity pattern, worker density and the dynamics of disease transmission in social insects. Journal of Theoretical Biology 226:45–51.

Pull, C. D., L. V. Ugelvig, F. Wiesenhofer, A. V. Grasse, S. Tragust, T. Schmitt, M. J. Brown, and S. Cremer. 2018. Destructive disinfection of infected brood prevents systemic disease spread in ant colonies. Elife 7:e32073.

Richard, F. J., A. Aubert, and C. M. Grozinger. 2008. Modulation of social interactions by immune stimulation in honey bee, Apis mellifera, workers. BMC Biology 6:50.

Richard, F.-J., H. L. Holt, and C. M. Grozinger. 2012. Effects of immunostimulation on social behavior, chemical communication and genome-wide gene expression in honey bee workers (apis mellifera). BMC Genomics 13:558.

Rueppell, O., M. K. Hayworth, and N. Ross. 2010. Altruistic self-removal of health-compromised honey bee workers from their hive. Journal of Evolutionary Biology 23:1538–1546.

Simone-Finstrom, M., and M. Spivak. 2010. Propolis and bee health: the natural history and significance of resin use by honey bees. Apidologie 41:295–311.

Smith, A. A., B. Hölldobler, and J. Liebig. 2012. Queen-specific signals and worker punishment in the ant Aphaenogaster cockerelli: the role of the Dufour’s gland. Animal Behaviour 83:587–593.

Spivak, M. 1996. Honey bee hygienic behavior and defense against Varroa jacobsoni. Apidologie 27:245–260.

Stow, A., D. Briscoe, M. Gillings, M. Holley, S. Smith, R. Leys, T. Silberbauer, C. Turnbull, and A. Beattie. 2007. Antimicrobial defences increase with sociality in bees. Biology Letters 3:422–424.

Sullivan, K., E. Fairn, and S. A. Adamo. 2016. Sickness behaviour in the cricket Gryllus texensis: comparison with animals across phyla. Behavioural Processes 128:134–143.

Turnbull, C., S. Hoggard, M. Gillings, C. Palmer, A. Stow, D. Beattie, D. Briscoe, S. Smith, P. Wilson, and A. Beattie. 2011. Antimicrobial strength increases with group size: implications for social evolution. Biology Letters 7:249–252.

van Zweden, J. S., and P. d’Ettorre. 2010. Nestmate recognition in social insects and the role of hydrocarbons. In G. J. Blomquist and A.-G. Bagnères, eds., Insect Hydrocarbons: Biology, Biochemistry and Chemical Ecology. Cambridge University Press, Cambridge.

